# Ethanol stress stimulates sumoylation of transcription factor Cst6 which restricts expression of its target genes

**DOI:** 10.1101/758607

**Authors:** Veroni S. Sri Theivakadadcham, Emanuel Rosonina

**Affiliations:** Department of Biology, York University, Toronto, Ontario, Canada

**Author notes:** Address correspondence to Emanuel Rosonina.

## Abstract

Sumoylation is an essential post-translational modification that functions in multiple cellular processes, including transcriptional regulation. Indeed, transcription factors represent one of the largest groups of proteins that are modified by the SUMO peptide. Multiple roles have been identified for sumoylation of transcription factors, including regulation of their activity, interaction with chromatin, and binding site selection. Here, we examine how Cst6, a bZIP-containing sequence-specific transcription factor in *Saccharomyces cerevisiae*, is regulated by sumoylation. Cst6 is required for survival during ethanol stress and has roles in the utilization of carbon sources other than glucose. We find that Cst6 is sumoylated to appreciable levels in normally growing yeast at Lys residues 139, 461 and 547, and that its sumoylation level increases in ethanol and oxidative stress conditions, but decreases if ethanol is used as the sole carbon source. To understand the role of Cst6 sumoylation during ethanol stress, we generated a yeast strain that expresses a non-sumoylatable mutant form of Cst6. Cellular levels of the mutant protein are moderately reduced compared to the wild-type form, implying that sumoylation promotes Cst6 stability. Although the mutant can bind DNA, chromatin immunoprecipitation (ChIP) analysis shows that its occupancy level is significantly reduced on promoters of some ethanol stress-regulated genes, suggesting that Cst6 recruitment is attenuated or delayed if it can not be sumoylated. Furthermore, impaired Cst6 sumoylation in the mutant strain correlates with elevated expression of some target genes, either constitutively or during induction by ethanol stress. This is most striking for *RPS3*, which shows dramatically increased expression in the mutant strain. Together, our results suggest that sumoylation controls multiple properties of Cst6 to limit the expression of its target genes.

## INTRODUCTION

Transcription factors maintain cellular functions by integrating external signal information into gene expression programs that are transmitted through signalling pathways. Often, these signalling pathways orchestrate the function of transcription factors through post-translational modifications (PTMs) including acetylation, phosphorylation, and ubiquitination (Carr et al. 2015). In recent years, regulation of transcription factors by sumoylation has gained increased attention. Sumoylation, a reversible post-translational modification that plays an essential role in many cellular processes, involves the covalent attachment of a ~12 kDa SUMO (*S*mall *U*biquitin-like *Mo*difier) peptide to specific lysine residues of substrate proteins (Cubeñas-Potts and Matunis 2013; Makhnevych et al. 2009). Similar to ubiquitination, sumoylation is a dynamic process that requires catalysis by three classes of enzymes: activation of SUMO by an E1 enzyme, conjugation to target proteins by an E2 enzyme (Ubc9), and facilitation of transfer by E3 ligases (Flotho and Melchior 2013). In contrast to ubiquitination, in which E3 ubiquitin ligases are the primary determinants of substrate specificity, Ubc9 can directly specify its targets via the SUMO site consensus sequence ΨKxD/E, where Ψ is a hydrophobic residue, K is the lysine to be modified, x is any amino acid, and D/E represents an acidic residue (Dye and Schulman 2007). Sumoylation is a reversible modification, in which SUMO can be removed from substrate proteins by SUMO-specific isopeptidases (SUMO proteases), including a family of sentrin/SUMO-specific proteases (SENPs) in mammals and the Ubl- specific proteases (Ulp1 and Ulp2) in budding yeast (Kunz et al. 2018; Flotho and Melchior 2013).

Proteomics studies, in which SUMO-conjugated proteins were isolated and identified through mass spectrometry, have determined that within the sumoylome of yeast and mammalian cells, proteins involved in transcription are among the largest classes of SUMO targets (Hendriks et al. 2014; Hendriks and Vertegaal 2016; Esteras et al. 2017; Hannich et al. 2005). This includes transcription factors, RNA polymerase II, transcriptional co-regulators and a variety of chromatin regulatory factors. Among these, sequence specific DNA-binding transcription factors represent the largest group of SUMO conjugates (Rosonina et al. 2017). In most cases, sumoylation of these transcription factors has been found to associate with transcriptional repression or deactivation. This can be achieved by recruiting various repressor complexes to chromatin, as for transcription factors NFAT, Elk-1 and MafG (Motohashi et al. 2006; Yang and Sharrocks 2004; Nayak et al. 2009), or by facilitating their clearance from promoters of induced genes, which is the case for transcription factors Gcn4 and c-Fos (Tempe et al. 2013; Rosonina et al. 2012). Genome-wide chromatin immunoprecipitation (ChIP) analyses showed that sumoylation-deficient forms of three human transcription factors, MITF, glucocorticoid receptor, and androgen receptor, occupied far more genomic sites than their wild-type counterparts (Bertolotto et al. 2011; Paakinaho et al. 2014; Sutinen et al. 2014). Similarly, we recently demonstrated that the sumoylation of the budding yeast bZIP transcription factor Sko1 is required to prevent it from binding to numerous non-target promoters (Sri Theivakadadcham et al. 2019). Together, these observation have led to a model in which controlling binding-site specificity has been proposed as a conserved and general role for sumoylation in regulating transcription factors (Rosonina 2019).

Whereas the studies mentioned above demonstrate that sumoylation can promote the dissociation of transcription factors from DNA at inappropriate genomic sites or during deactivation, for some transcription factors, sumoylation has been shown to facilitate their association with DNA. For example, sumoylation enhances the DNA-binding ability of Pax-6 and CREB1, thereby positively regulating transcription of their target genes (Yan et al. 2010; Chen et al. 2014). Furthermore, sumoylated proteins are detected at promoters of transcriptionally active genes, including constitutive and activated inducible genes, indicating that transcription factor sumoylation can function during active transcription (Rosonina et al. 2010). The individual effects of transcription factor sumoylation, therefore, can vary with substrate and context, as sumoylation can function during the initial stages of transcription activation, during active transcription, during the deactivation process, or to promote transcriptional silencing.

To extend our understanding of transcription factor sumoylation, we focused on the budding yeast bZIP transcription factor, Cst6 (*C*hromosome *St*ability-*6*), which was identified as a potential SUMO target in large-scale proteomics studies (Wohlschlegel et al. 2004; Hannich et al. 2005). Cst6, also known as Aca2, is essential during ethanol stress and in the presence of non-optimal carbon sources (Saleem et al. 2010; Garcia-Gimeno and Struhl 2000). It contains a bZIP DNA binding domain, binds to DNA as a homodimer or a heterodimer with Aca1, another yeast bZIP transcription factor, and it recognises cAMP response element (CRE)-like promoter sequences (Garcia-Gimeno and Struhl 2000). A recent study showed that the Cst6 binding site (5’-GTGACGT-3’) has an additional guanine nucleotide at the 5’ end of the consensus CRE motif (5’-TGACGT-3’), implying that it has a somewhat different binding site preference than other bZIP transcription factors (Liu et al. 2016). Through genome-wide binding site analysis, the study also identified 59 protein-coding genes as putative targets for Cst6 regulation. Gene ontology (GO) analysis of these genes revealed that Cst6 controls different cellular processes including transcription, cellular respiration, gluconeogenesis, stress response, and pseudohyphal growth, signifying the importance of its function in budding yeast (Liu et al. 2016).

In the present study, we demonstrated that Cst6 is multi-sumoylated on three lysine residues (K139, K461, and K547), and that the level of its SUMO modification increases during ethanol and oxidative stress. Whereas the Cst6 sumoylation increases with increased duration and dose of ethanol stress, it is significantly reduced if ethanol is used as the sole carbon source in the medium. We further demonstrate that a mutant, non-sumoylatable form of Cst6 shows reduced or delayed association with its target promoters during ethanol stress. This might be partly explained by a moderate decrease in the abundance of Cst6 in the absence of its sumoylation. Finally, we find that the Cst6 mutant strain shows elevated expression of Cst6 target genes either constitutively, or during exposure to ethanol stress. Collectively, these results suggest that sumoylation enhances Cst6 binding to ethanol-induced target promoters to limit the transcription of its target genes.

## RESULTS

### Cst6 is sumoylated during normal growth

In large-scale proteomics studies, Cst6 was identified as a putative SUMO target (Wohlschlegel et al. 2004; Hannich et al. 2005). In order to confirm Cst6 sumoylation, we epitope tagged *CST6* with a 6xHA tag using homologous recombination in the common lab yeast strain W303a. The presence of the HA tags had no effect on cell growth as confirmed by a spot assay performed under several growth conditions (Figure S1A and B). Cst6-6HA was then immunoprecipitated (IPed) using an HA antibody from cell lysate, prepared in the presence of *N*-ethylmaleimide (NEM), an inhibitor of SUMO proteases, which was then analyzed by SUMO and HA immunoblotting. Supporting that Cst6 is sumoylated, bands were detected on the SUMO immunoblot for the Cst6-6HA IP (Figure 1A). As a positive control, Cst6-6HA sumoylation levels were compared alongside another bZIP transcription factor, Sko1, which is known to be polysumoylated during non-stress conditions (Sri Theivakadadcham et al. 2019). In addition to Sko1-6HA or Cst6-6HA-specific bands, a band of variable intensity was detected at ~250 kDa in the SUMO immunoblots of HA IPs (arrow in Figure 1A and throughout). Because it is also detected in control HA IPs from lysates lacking HA-fused proteins (Figure 1A and see Figure 4B), we believe that this band corresponds to a sumoylated protein that non-specifically co-purifies during native IPs. Like for Sko1, multiple SUMO-specific bands were detected for Cst6, including a major band and multiple fainter higher molecular weight bands, which is likely the result of multi- or polysumoylation (Figure 1A). To further validate that these bands are due to conjugation of SUMO with Cst6, lysates were prepared in the absence of NEM, in which the level of sumoylation is expected to be reduced significantly due to the uninhibited activity of SUMO proteases. Indeed, in the absence of NEM, Cst6 sumoylation is undetectable (Figure S1C, compare lanes 1 and 2). In Cst6-6HA IPs, two bands are observed on long-exposed HA immunoblots: an intense band slightly below 100 kDa, which corresponds to unmodified Cst6-6HA, and a fainter higher band that co-migrates with the major sumoylated form of Cst6 (~125 kDa) seen on the SUMO immunoblot (indicated by an asterisk on Figures 1A and S1C). Supporting that this band corresponds to sumoylated Cst6, it is not detected when the IP experiment is performed in the absence of NEM (Figure S1C).

**Figure 1:**
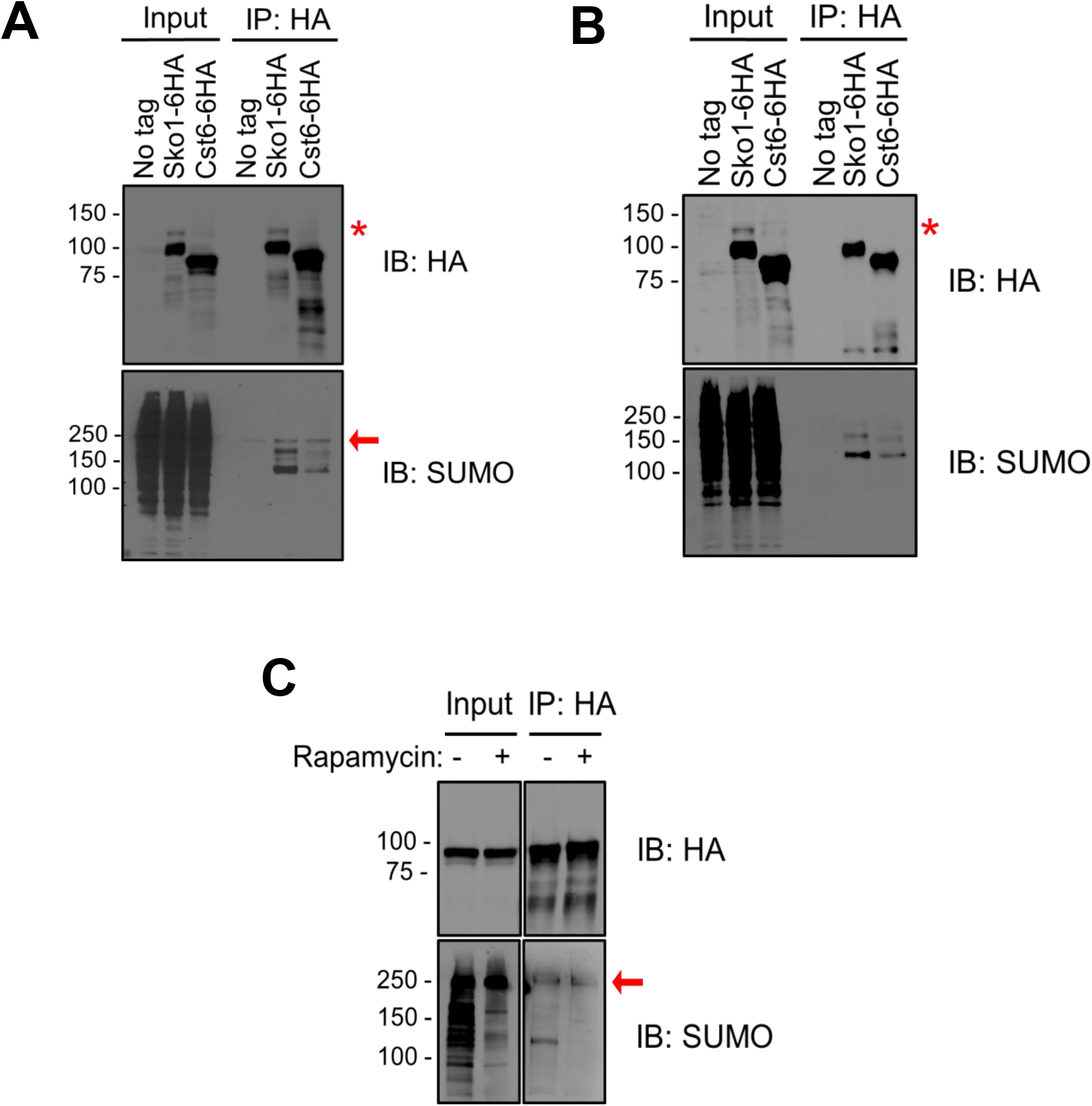
bZIP transcription factor Cst6 is sumoylated. **(A)** Detection of Cst6 sumoylation using immunoprecipitation (IP) with lysates prepared in non-denaturing conditions. 6HA-tagged Sko1 and Cst6 were IPed using HA-conjugated beads from cell lysates prepared in the presence of *N*-ethylmaleimide (NEM) and analyzed by immunoblot with HA or SUMO antibodies, as indicated. ‘No tag’ refers to a sample from the parental strain (W303a) expressing no HA-tagged proteins, used as a negative control. Sko1 is a known SUMO target (Sri Theivakadadcham et al. 2019) and was used as a positive control. **(B)** Sumoylation of Cst6 detected by IP from protein extracts prepared under denaturing conditions. Extracts were generated by TCA precipitation, treated with SDS, and boiled before IPing with HA conjugated beads. Extracts were analyzed by immunoblot with the antibodies indicated. **(C)** Cst6 sumoylation is abolished when Ubc9 is conditionally removed from the nucleus. Cst6-6HA was IPed from a Ubc9 anchor away strain using an HA antibody from lysates prepared from cells treated with or without rapamycin for 30 min. Sumoylation levels were determined by IP-immunoblot analysis, as in *A*. Red asterisk (*) indicates the position of SUMO-modified Cst6-6HA on HA blots. Arrow indicates the position a non-specific, co-purifying protein detected on SUMO blots.

Additional experiments were performed to confirm Cst6 sumoylation. To eliminate the possibility that bands observed in the SUMO blot are derived from sumoylated proteins that co-IPed with Cst6, lysates were prepared under denaturing conditions prior to IP. Cellular proteins were extracted with trichloroacetic acid (TCA), treated with sodium dodecyl sulfate (SDS), and boiled before IPing with HA-conjugated beads. Under these conditions, Cst6 sumoylation was still detected, as observed in the SUMO and HA blots in Figure 1B, which indicates that the signal observed in the SUMO immunoblot is due to SUMO-conjugated Cst6. As a final method to confirm Cst6 sumoylation, we used a strain in which Ubc9, the sole SUMO conjugating enzyme, is fused to an “anchor away” tag that causes its translocation from the nucleus to the cytoplasm when cells are exposed to rapamycin (Haruki et al. 2008), thereby conditionally blocking nuclear sumoylation events. In the presence of rapamycin, the level of global sumoylation is dramatically reduced, as seen in the SUMO blot of denatured lysate inputs (Figure 1C). Cst6-6HA was IPed using an HA antibody from lysates of cultures treated with or without rapamycin and analyzed by HA and SUMO immunoblots. Sumoylation of Cst6 was abolished after rapamycin treatment, further confirming that the transcription factor Cst6 is a target for sumoylation in normally growing yeast (Figure 1C).

### Cst6 sumoylation increases during oxidative and ethanol stress conditions

Numerous studies have reported the importance of sumoylation in maintaining cell homeostasis during environmental stress (e.g. (Enserink 2015; Niskanen et al. 2015)). Cst6 is part of the stress-responsive transcriptional regulatory network and it was previously shown to be involved in regulating transcription during ethanol, oxidative and heat shock stress (Suzuki et al. 2011; Wu and Li 2008; Liu et al. 2016; Garcia-Gimeno and Struhl 2000). To explore the effect of stress on Cst6 sumoylation, we examined its sumoylation levels by IP-immunoblot in lysates derived from cells grown under different stress conditions (Figures 2A and B). Cst6 sumoylation is significantly increased during ethanol and oxidative stress conditions and reduced during osmotic stress, while no change was observed during amino acid starvation or heat shock. Cst6 sumoylation during ethanol stress was further confirmed with additional IP experiments in which NEM was included or excluded. As shown in Figure S1C, Cst6 sumoylation increases with ethanol stress in samples prepared in the presence of NEM and it is abolished in its absence (compare lanes 1, 3 and 4). Our data demonstrates that sumoylation of Cst6 increases during ethanol and oxidative stress, and further experiments were performed to investigate the role that sumoylation plays in regulating Cst6 function in these conditions.

**Figure 2:**
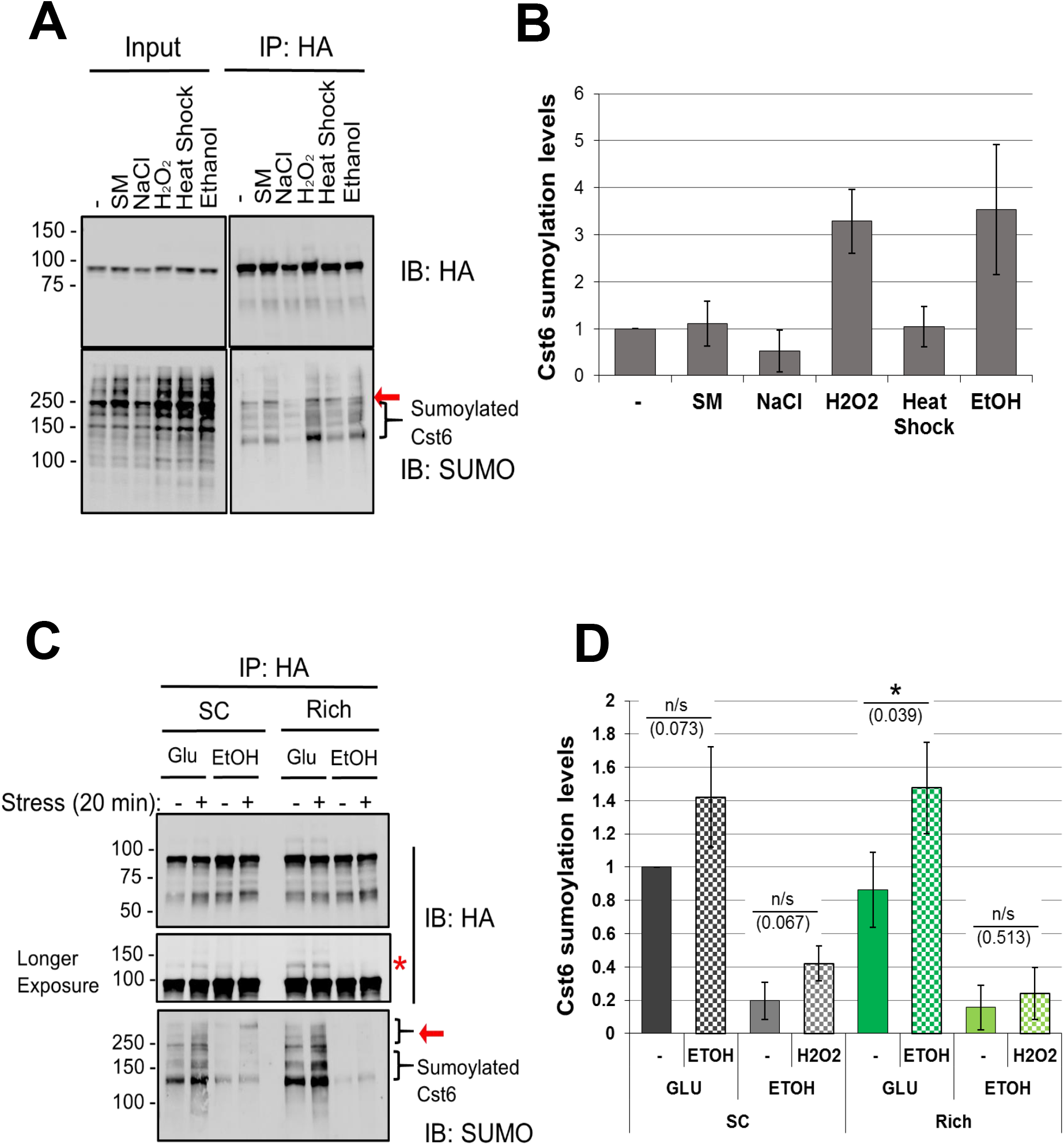
Sumoylation level of Cst6 is elevated during oxidative and ethanol stress. **(A)** Cells expressing Cst6-6HA were treated with the indicated stressors for 20 min and Cst6 sumoylation levels were determined by IP-immunoblot, as in Figure 1A. The following stress conditions were used: 0.5 μg/mL of sulfometuron methyl (SM), which induces amino acid starvation; 0.4 M NaCl for osmotic stress; 100 mM H_2_O_2_ for oxidative stress; 37°C for heat shock; and 5% ethanol (EtOH) for ethanol stress. **(B)** Quantification of Cst6 sumoylation levels for analysis shown in *A*. On the SUMO immunoblot, the intensity of signals below the non-specific bands were quantified using ImageJ, normalized to the HA IP signal intensities, then plotted relative to the first sample. **(C)** Cells were grown in SC or rich media containing either 2% glucose (Glu) or 1% ethanol as a carbon source, then treated with or without stress (100 mM H_2_O_2_ for cells grown in ethanol-containing medium or 5% ethanol for cells grown in glucose-containing medium) for 20 min. Sumoylation levels were determined by IP-immunoblot as in Figure 1A. **(D)** Quantification of Cst6 sumoylation levels for analysis shown in *C*. Red asterisk (*) indicates the position of SUMO-modified Cst6 on HA immunoblot. Arrows refer to a non-specific band on the SUMO blots. Error bars represent standard deviation of three independent experiments. Significant differences between the non-stress and the stress samples under each media conditions were calculated using Student’s *t*-test (*, *P* < 0.05; n/s, not significant).

Previous studies on Cst6 highlighted its importance for cell growth in the presence of alternative carbon sources (Garcia-Gimeno and Struhl 2000; Liu et al. 2016). Typically, lab yeast strains are grown in media containing glucose, but other carbon sources, such as glycerol, ethanol, and raffinose, can be used as alternatives. Ethanol, therefore, can serve as a carbon source, or as a stress condition when used at high concentrations in the presence of glucose as a carbon source. In the absence of Cst6, cells grow poorly in alternative carbon sources, including ethanol (Garcia-Gimeno and Struhl 2000; Liu et al. 2016). We wished to investigate whether the level of Cst6 sumoylation changes with different types of media and carbon sources used for growth. To test this, we grew the Cst6-6HA strain in SC (Synthetic Complete) or rich medium that was supplemented with either 2% glucose or 1% ethanol as a carbon source, and cells were then treated with or without a stress condition, as indicated on Figure 2C. Cells were lysed and Cst6-6HA was then IPed and analyzed by HA and SUMO immunoblots. Strikingly, the level of Cst6 sumoylation was significantly lower when grown in ethanol versus glucose (Figures 2C and D), which might reflect the lower levels of global sumoylation observed when ethanol is the sole carbon source (Figure S2A). As expected, Cst6 sumoylation levels were significantly increased when cells grown in rich, glucose-containing medium were exposed to ethanol stress, but oxidative stress did not show a statistically significant effect on yeast grown in ethanol-containing medium (Figures 2C and D). In summary, normally growing yeast display a basal level of Cst6 sumoylation that can be increased by ethanol or osmotic stress, or decreased if ethanol is used as the sole carbon source. Based on these results, we proceeded to investigate what role sumoylation plays in regulating Cst6 function during non-stress and ethanol stress conditions using rich, glucose-containing medium.

To determine whether the effect of ethanol stress on Cst6 sumoylation levels is dose-dependent, we treated the Cst6-6HA strain with different concentrations of ethanol for 20 mins. Cells were then lysed, IPed and analyzed by HA and SUMO immunoblots. Indeed, Cst6 sumoylation was elevated with increased concentrations of ethanol (Figures 3A and B). This might at least partly be due to an increase in global sumoylation that occurs with exposure to up to 7.5% ethanol (Figures S2B and C). Next, we examined whether Cst6 sumoylation is dependent on the exposure time for treatment with ethanol. Sumoylation levels were determined from Cst6-6HA-expressing cells treated with 5% ethanol for a range of times. As shown in Figures 3C and D, Cst6 sumoylation levels increased with prolonged exposure to ethanol stress. Based on these results, and conditions commonly used by others, further ethanol stress treatments were performed at 5% for 20 min.

**Figure 3:**
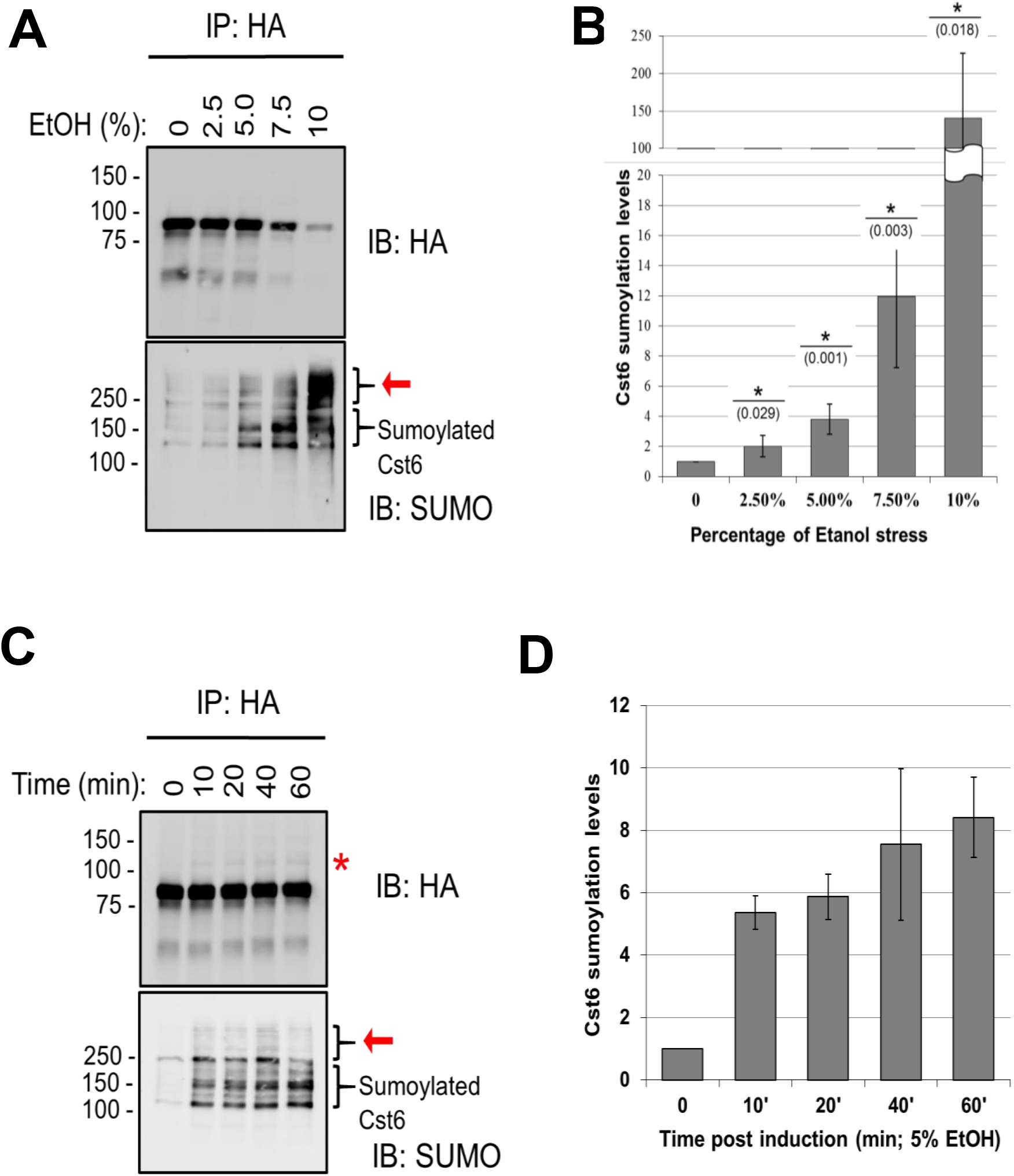
Cst6 sumoylation level during ethanol stress is dependent on the dose and duration of stress exposure. **(A)** Cst6 sumoylation is elevated with increasing concentrations of ethanol exposure. Cells expressing Cst6-6HA were treated with the indicated concentrations of ethanol for 20 mins and then sumoylation levels were determined as in Figure 1A. **(B)** Quantification analysis of Cst6 sumoylation levels for *A* as in Figure 2B. **(C)** Cst6 sumoylation is increased with prolonged ethanol stress. 6HA-tagged cells were treated with 5% ethanol for indicated time points and then the sumoylation levels were characterized as in Figure 1A. **(D)** Quantification analysis of Cst6 sumoylation levels for *C* as in Figure 2B. Red asterisk (*) indicates SUMO modified Cst6 on HA immunoblot. Arrows indicate non-specific bands on SUMO blots. Error bars represent standard deviation of three independent experiments. Significant differences between the non-stress sample and with each stress conditions were calculated using Student’s *t*-test (*, *P* < 0.05; n/s, not significant).

### Cst6 is sumoylated on lysine residues 139, 461 and 547

Using SUMO site prediction software (GPS-SUMO, SUMOplot, and JASSA), we identified three putative SUMO sites for Cst6: K139, K461, and K547, all of which fall within SUMO site consensus motifs (Figure 4A) (Zhao et al. 2014; Beauclair et al. 2015). K139 is situated just upstream of the Aca-specific region (a region identical in Aca1 and Aca2/Cst6 (Garcia-Gimeno and Struhl 2000)), K461 is found within the bZIP domain, and K547 is at the C-terminus of Cst6. To determine whether these are actual sumoylation sites, we generated yeast strains expressing Lys-to-Arg substitution mutants of all three putative SUMO sites, since Arg cannot be sumoylated (strains Cst6-K139R-6HA, Cst6-K461R-6HA and Cst6-K547R-6HA). We used IP-immunoblot to determine sumoylation levels and found that the individual K139R or K547R mutations decreased Cst6 sumoylation more than the K461R mutation, but they did not completely abolish Cst6 sumoylation (Figures 4B and C). Therefore, we generated double-(“D.MT”; Cst6-K461,547R-6HA) and triple-mutant (“T.MT”; Cst6-K139,461,547R-6HA) strains to assess their effects on Cst6 sumoylation. The D.MT mutations partially reduced Cst6 sumoylation, whereas the T.MT mutations abolished sumoylation of Cst6 under normal and ethanol-stress conditions (Figures 4B, C, and S1C, lanes 5-8). These results strongly suggest that Cst6 is sumoylated at all three lysines and, therefore, Cst6 is multi-sumoylated. The T.MT strain (hereafter referred to as Cst6 MT), alongside the CST6-6HA strain (hereafter referred to as Cst6 WT), were then used to explore roles for Cst6 sumoylation.

**Figure 4:**
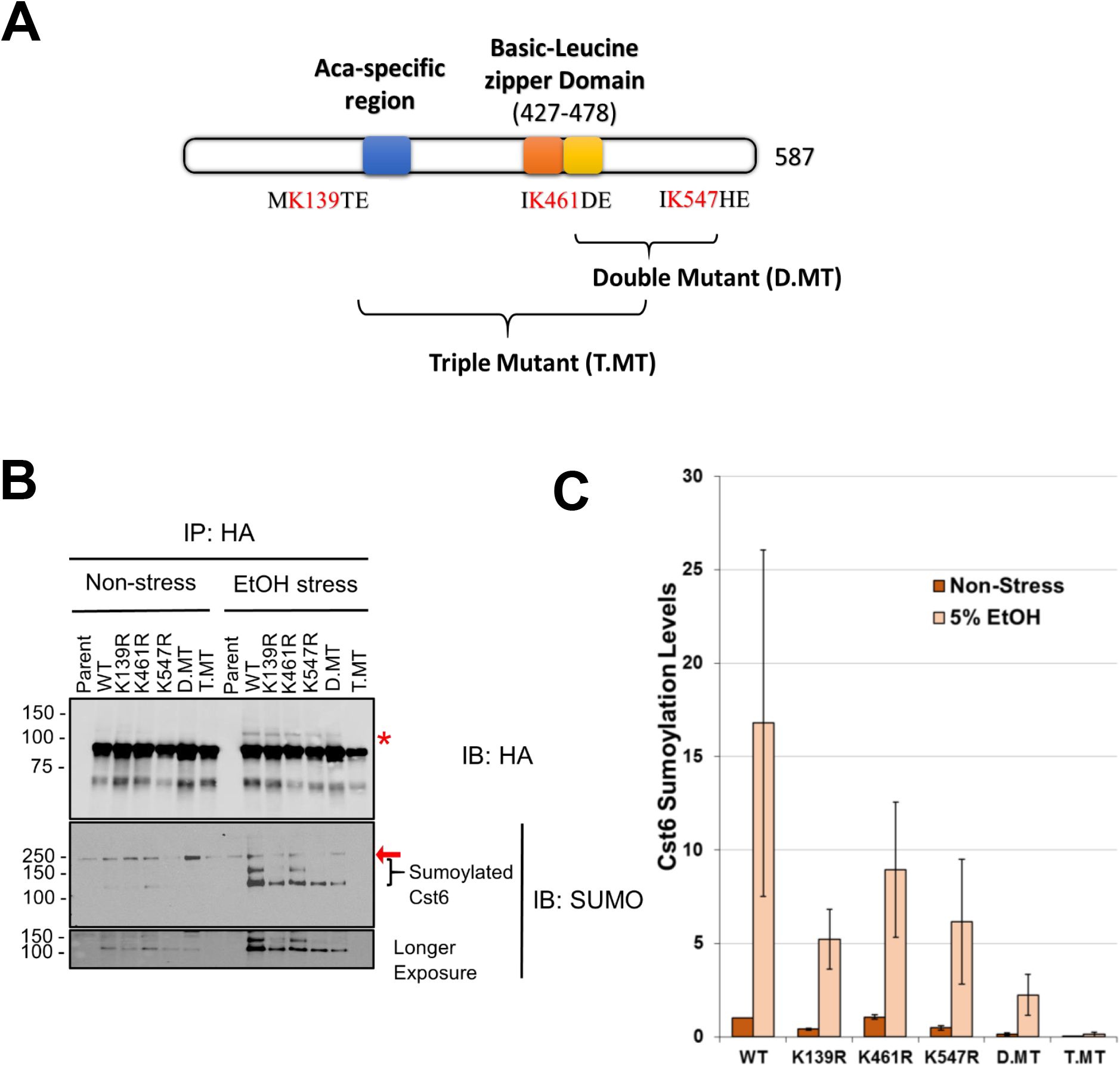
Cst6 is multi-sumoylated at K139, K461 and K547 during non-stress and ethanol stress conditions. **(A)** Schematic diagram of Cst6 indicating the Aca-specific region and basic leucine Zipper (bZIP) domain. Putative SUMO sites on Cst6 were identified as lysine residues 139, 461 and 547 using SUMO site prediction software and are indicated with the encompassing SUMO consensus motifs. **(B)** Cst6 is sumoylated on all three putative SUMO sites during non-stress and ethanol stress conditions. Using site-directed Lys-to-Arg mutagenesis, SUMO-site mutant strains of Cst6 [Cst6-K139R, Cst6-K461R, Cst6-K547R, Cst6-K461,547R (D.MT), and Cst6-K139,461,547R (T.MT)] were generated. Sumoylation levels of “No tag”, Cst6-6HA (WT), and the mutants were characterized as in Figure 1A. **(C)** Quantification of Cst6 sumoylation levels for *B* as in Figure 2B. Red asterisk (*) indicates the SUMO modified Cst6 on HA blot. Arrows indicate non-specific bands on SUMO blots. Error bars represent standard deviation of two independent experiments.

### Sumoylation of Cst6 is not essential for cell viability but plays a minor role in its stability

To study whether sumoylation of Cst6 has a role in cell fitness, a spot assay was conducted. Supporting previous studies, cells lacking Cst6 showed a growth defect during ethanol stress, when glucose was the carbon source (Figure 5A; (Garcia-Gimeno and Struhl 2000; Liu et al. 2016)). However, under either non-stress conditions or when exposed to different concentrations of ethanol, no growth defect was observed for Cst6 MT, implying that sumoylation of Cst6 is not essential for cell viability or survival during ethanol stress (Figure 5A). In addition, we compared growth levels on medium in which ethanol (either 1% or 3%) was the carbon source. Deletion of *CST6* led to severe growth defects, but the Cst6 MT strain once again grew as well as the Cst6 WT strain, and addition of hydrogen peroxide, to induce oxidative stress, had no effect (Figure 5B). These results together indicate that sumoylation of Cst6 is not essential for cell viability or fitness under any of the tested growth conditions.

**Figure 5:**
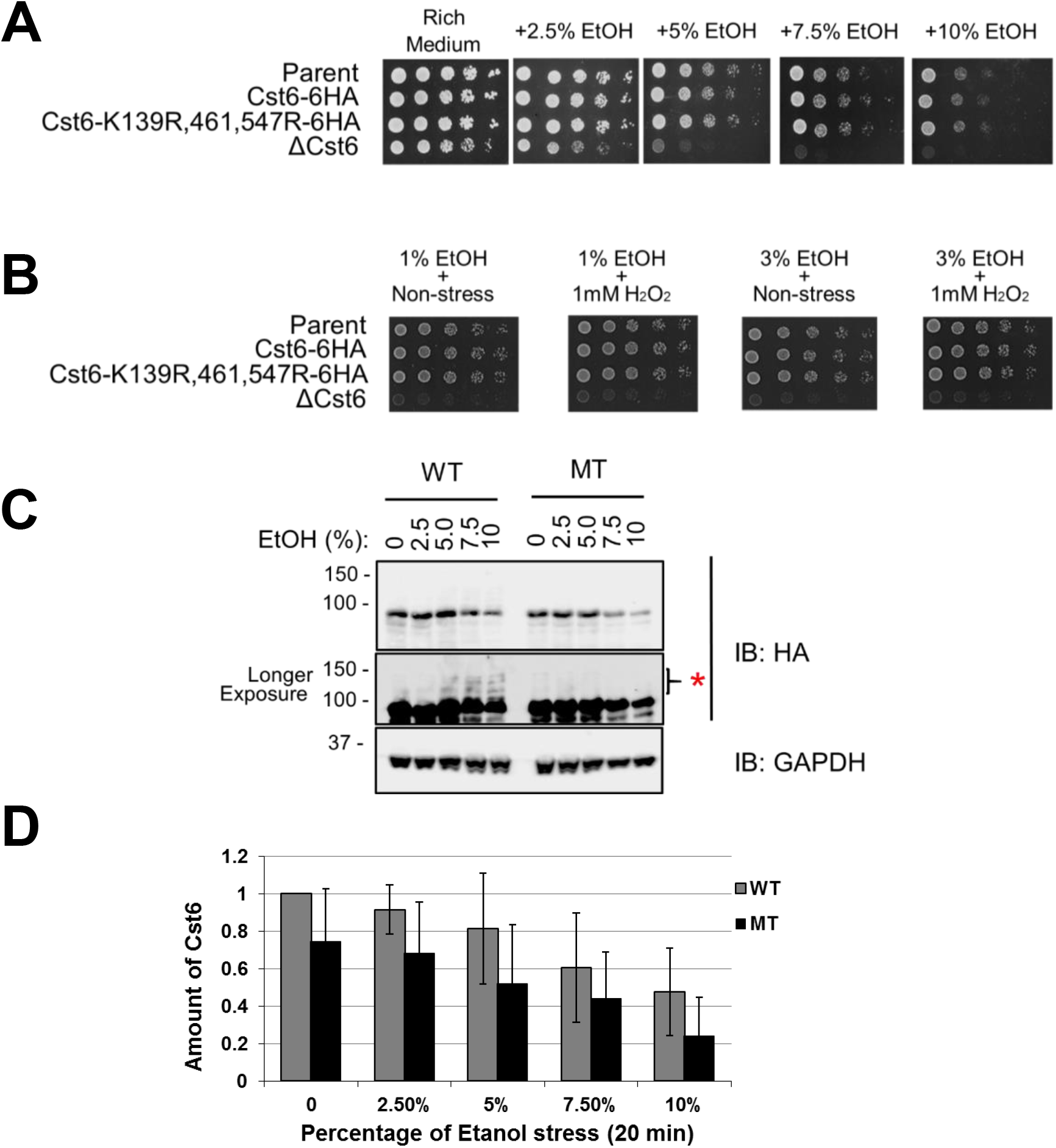
Sumoylation of Cst6 is not essential for cell viability but influences its abundance. **(A)** Cst6 sumoylation is not essential for cell viability during ethanol stress on medium containing 2% glucose. Cell growth fitness was assessed on glucose-containing rich-medium plates, under non-stress and indicated ethanol stress conditions, using the spot assay technique with a five-fold serial dilution series. Plates were then incubated for two days at 30°C. **(B)** Cst6 sumoylation is not essential for cell viability when ethanol is the carbon source. Cell growth was assessed on ethanol-containing rich-medium plates under non-stress and oxidative stress conditions, by spot assay. Plates were incubated for two days at 30°C. **(C)** Cst6 abundance is reduced when its sumoylation sites are mutated. Lysates (from cells expressing Cst6-6HA) treated with the indicated percentages of ethanol for 20 min were analyzed by HA and GAPDH immunoblots. **(D)** Quantification analysis of Cst6 levels for *C*. Intensity of HA signals were normalized to corresponding GAPDH signals, then plotted relative to the first sample. Red asterisk (*) indicates the SUMO modified form of Cst6 on the HA blot. Error bars represent standard deviation of three independent experiments.

Next, we examined whether ethanol stress or sumoylation affect Cst6 protein abundance. Cell lysates were prepared from cells treated with different concentrations of ethanol for both WT and MT strains and we analyzed Cst6 levels by HA immunoblot. Increasing ethanol concentration correlated with decreased abundance of Cst6, suggesting that Cst6 is degraded during ethanol stress (Figures 5C and D). Interestingly, there appears to be an inverse correlation between the level of Cst6 sumoylation and its abundance (see also Figures 3A and B). In conditions where sumoylation of Cst6 increases, protein levels of Cst6 are reduced, suggesting that sumoylation is involved in Cst6 protein stability. To explore this, we compared Cst6 protein levels in the WT and MT strain after exposure to increasing levels of ethanol. As shown in Figures 5C and D, Cst6 MT also becomes less abundant with increased ethanol stress, but the level of Cst6 MT is consistently less than the level of Cst6 WT, ranging from ~25% less in the absence of ethanol to ~49% less in the presence of 10% ethanol. Whereas sumoylation is not needed for the ethanol-dependent reduction in Cst6 protein levels, our data suggests that Cst6 sumoylation plays a role in enhancing its stability in normal and ethanol stress conditions.

### Sumoylation enhances Cst6 promoter binding and restricts expression of its target genes

Sumoylation of bZIP transcription factors can function in both promoter clearance (e.g. c-Fos, Gcn4 and Sko1; (Rosonina et al. 2012; Tempe et al. 2013; Sri Theivakadadcham et al. 2019)) and enhancing DNA binding (e.g. CREB1; (Chen et al. 2014)). Therefore, it is possible that sumoylation of Cst6 plays a role in regulating its association or dissociation with promoters of its target genes. To investigate this, chromatin immunoprecipitation (ChIP) was used to compare the occupancy levels of Cst6 WT and Cst6 MT on ethanol-induced target promoters. The occupancy levels of Cst6-WT significantly increased during ethanol stress on all the target genes tested, except on the control gene *PMA1,* which is not a target of Cst6 (Figure 6A). Cst6 MT was also recruited to Cst6 target genes during ethanol stress but showed significantly reduced occupancy compared to WT after 15 min of ethanol stress at three of the tested genes (*PYC1*, *YAP6*, and *RPS3*). Although efforts were made to match conditions and timing between replicate experiments, there were significant deviations in Cst6 occupancy levels among the replicates, which hampered determination of statistical significance between Cst6 WT and MT occupancy for other genes and time points. Nonetheless, comparing the occupancy patterns of WT and MT suggests that recruitment of non-sumoylatable Cst6 is generally delayed and/or attenuated on target genes, which implies that sumoylation is important for the timely association of Cst6 with target promoters during ethanol stress.

**Figure 6:**
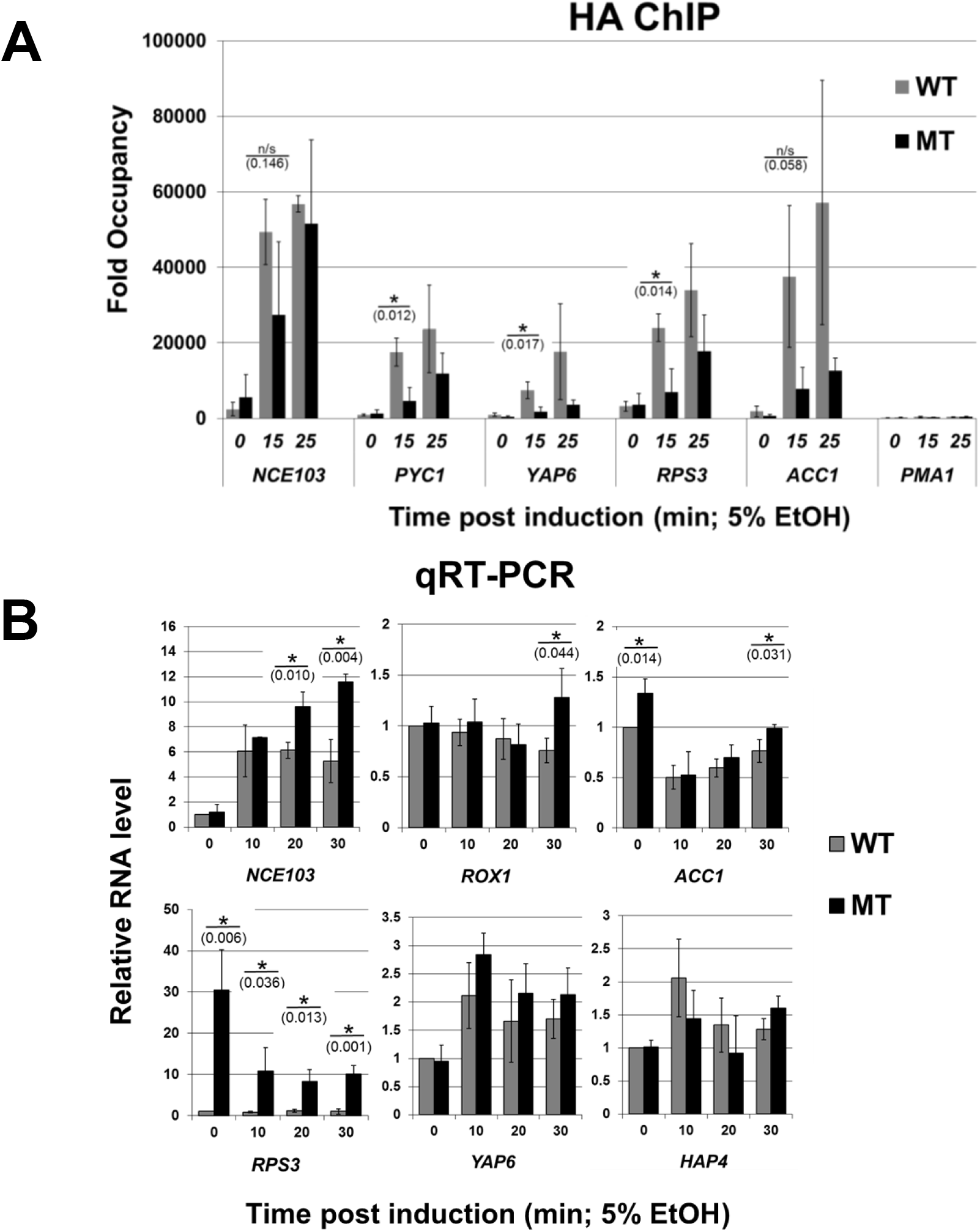
Sumoylation enhances Cst6 promoter binding and restricts expression of some target genes. **(A)** Quantitative HA ChIP analysis of Cst6 WT and Cst6 MT occupancy levels on its ethanol stress-regulated target gene promoters after induction with 5% ethanol for indicated times. Gene occupancy signals were normalized to an internal control, an untranscribed region of Chromosome V. **(B)** Quantitative RT-PCR analysis was performed on RNA isolated from *CST6 WT* and *Cst6 MT* strains. Cells were treated with 5% ethanol and harvested after the indicated time points. mRNA levels were normalized to levels of the *25S* rRNA. Error bars represent standard deviation of three replicates. Significant differences between Cst6 WT and Cst6 MT under each tested condition were determined by Student’s *t*-test (*, *P* < 0.05).

To further study the effect of Cst6 sumoylation on gene expression, we investigated how sumoylation might regulate transcription of Cst6 target genes. The majority of initial studies on roles for sumoylation in regulating transcription factors pointed to roles for the modification in transcription silencing (Rosonina et al. 2012; Tempe et al. 2013; Motohashi et al. 2006; Nayak et al. 2009; Yang and Sharrocks 2004), whereas other studies then demonstrated roles for SUMO modification in transcriptional activation (e.g. (Yan et al. 2010; Chen et al. 2014)). To determine whether sumoylation of Cst6 is important for repression or activation of target genes, quantitative RT-PCR (qRT-PCR) analysis was performed on RNA isolated from the Cst6 WT and Cst6 MT strains. As seen in Figure 6B, the expression of various Cst6 target genes differs during ethanol stress. For example, although transcription of *NCE103, YAP6,* and *HAP4* was induced, the transcription of *ACC1* and *RPS3* was reduced, and no changes were observed for *ROX1.* In the Cst6 MT strain, expression patterns for most genes paralleled the patterns seen for Cst6 WT, with notable exceptions. Some genes showed elevated mRNA levels during later time points after exposure to ethanol stress (*NCE103, ROX1* and *ACC1*), some showed elevated levels during non-stress conditions (*ACC1* and *RPS3*), and some did not show significant differences (*YAP6,* and *HAP4*) (Figure 6B). Strikingly, *RPS3* showed dramatically elevated mRNA levels at all the indicated time points, suggesting that Cst6 sumoylation plays a major role in repressing expression of this gene. Overall, these results indicate that blocking Cst6 sumoylation affects expression of target genes and suggests that Cst6 sumoylation has a general repressive effect that is observed differentially at different targets.

## DISCUSSION

Extending our previous work on Gcn4 and Sko1, we have now shown that another bZIP transcription factor, Cst6, is also regulated by sumoylation in budding yeast (Rosonina et al. 2012; Sri Theivakadadcham et al. 2019). Gcn4 expression (and consequently, its sumoylation) is dependent on amino acid starvation, but like Sko1, a fraction of Cst6 molecules is sumoylated during normal growth conditions, suggesting that the modification regulates properties of the transcription factor not related to stress. However, Cst6 sumoylation increases with exposure to stress, specifically ethanol and oxidative stress, which we did not observe for Sko1.

Environmental stressors such as osmotic stress, oxidative stress, heat shock and ethanol stress lead to a dramatic increase in the level of protein sumoylation in mammalian, yeast and plant cells, but it is not fully understood how elevated sumoylation of specific target proteins affects their stress-related functions (Castro et al. 2012; Golebiowski et al. 2009; Lewicki et al. 2015; Chymkowitch et al. 2015). As the level of Cst6 sumoylation during ethanol stress was dependent on the dose and the duration of the stress, this provided us with an opportunity to examine how increased sumoylation might regulate properties of Cst6 during stress. Indeed, our analysis indicates that elevated sumoylation after ethanol stress likely promotes Cst6 recruitment or accumulation at target gene promoters and ensures that expression of its targets is not excessive.

Controlling the association of transcription factors with chromatin is a frequently reported function for sumoylation (Rosonina et al. 2017). For some transcription factors, sumoylation was shown to occur specifically after they bind to target promoters and the modification facilitates their subsequent removal from DNA, thereby enabling gene deactivation or restricting expression levels (e.g. Gcn4, Ikaros, and Sko1; (Rosonina et al. 2012; Apostolov et al. 2016; Sri Theivakadadcham et al. 2019)). As mentioned above, however, in other cases, sumoylation promotes the interaction of transcription factors with DNA (e.g. CREB1 and Pax6 (Yan et al. 2010; Chen et al. 2014)). As our ChIP analyses demonstrate that a sumoylation-deficient form of Cst6 can be recruited to DNA, sumoylation is not required for Cst6 to bind to its target promoters. However, our results support a role for sumoylation in enhancing the association of Cst6 with its binding sites. For one, as Cst6 sumoylation increases during exposure to ethanol stress, the levels of DNA-bound Cst6 also increase. Secondly, sumoylation-blocking mutations in Cst6 result in reduced chromatin occupancy at some target promoters.

As one possible mechanism for promoting the association of Cst6 with chromatin, sumoylation might increase its affinity for DNA through conformational changes or by promoting homodimerization or interactions with other proximal DNA binding transcriptional factors. In support of this, one of the Cst6 sumoylation sites that we identified, K461, is predicted to lie adjacent to the hydrophobic face of the leucine zipper region of its bZIP domain, which enables dimerization (Llorca et al. 2014). Phosphorylation of an analogous position of the leucine zipper of C/EBPβ is believed to enhance homodimerization, but more analysis is needed to explore how addition of a SUMO moiety at this residue affects pairing of Cst6 units, or heterodimerization with its paralog, Aca1 (Lee et al. 2010; Garcia-Gimeno and Struhl 2000). In an alternative mechanism for promoting increased Cst6 occupancy on DNA, sumoylation might enhance Cst6 stability, as has been shown for transcription factors Delta-Lactoferrin and NPHP7 (Escobar-Ramirez et al. 2015; Ramachandran et al. 2015). Indeed, our finding that Cst6 protein levels are reduced when its sumoylation sites are mutated supports this idea. In any case, whether through an active role in promoting its interaction with DNA, or through a less direct role in inhibiting its degradation, our study points to an important function for sumoylation in ensuring that high levels of Cst6 occupy its target gene promoters rapidly after exposure to ethanol stress.

Although Cst6 binds dozens of gene promoters in the presence of ethanol, the effect of Cst6 binding on their expression is gene dependent. For example, during ethanol stress, compared to wild-type cells, *NCE103* expression is dramatically reduced while *RPS3* expression is elevated in cells lacking Cst6 (Liu et al. 2016). Our analysis demonstrates, however, that impairing Cst6 sumoylation has a consistent effect on elevating transcription levels, wherever an effect was observed. This might reflect reduced occupancy levels for the sumoylation-deficient Cst6 mutant, suggesting that appropriate levels of Cst6 are normally needed to prevent excessive expression of targets. However, in normal conditions (i.e. time zero for the ethanol stress time course), we detected approximately equal levels of Cst6 WT and MT on the *RPS3* promoter, but expression of the gene was dramatically derepressed in a strain in which Cst6 cannot be sumoylated (Figure 6A and B). This points to a more direct role for sumoylation of Cst6 in inhibiting transcription. As with multiple metazoan transcription factors that are sumoylated, SUMO can directly impair transcription by promoting interactions with transcriptional corepressors, particularly histone deacetylase complexes (Ouyang and Gill 2009; Rosonina et al. 2017). Future studies will be key for understanding how sumoylation affects the molecular properties of Cst6, including its dimerization and interactions with other proteins and DNA, that lead to its transcriptionally repressive effects. Taken together, our study on Cst6 sumoylation provides further evidence that the modification controls the association of transcription factors with chromatin and fine-tunes transcription levels of target genes.

## MATERIALS AND METHODS

### Yeast strains

All *Saccharomyces cerevisiae* strains used in this study are listed in Table S1. Cst6 was epitope-tagged with a 6xHA tag (along with a *K. lactis TRP1* marker gene) using homologous recombination, as previously described by Knop et al. (1999). PCR-based point mutagenesis was used to generate strains expressing mutant forms of Cst6-HA. The Cst6-deleted strain was constructed using a *KanMX* deletion cassette, and all strains are derived from the W303a background strain.

### Media and cultivation

Yeast cultures were grown overnight in either rich (1% yeast extract, 2% peptone and 2% glucose) or Synthetic Complete (SC; 0.17% YNB, and 0.5% ammonium sulfate) medium (containing either 2% glucose or 1% ethanol as carbon source, or as otherwise noted) at 30°C and diluted to an optical density (OD_595nm_) of 0.2. Strains were then cultivated on a rotary shaker at 200 rpm until the OD reached 0.5-0.7 at which point, where appropriate, stressors were added directly to the culture as indicated in the figure captions or legends. Cells were then harvested by centrifugation at 3000 *g* for 5 min, followed by a wash with an experiment-specific buffer. For growth assays on agar plates (“spot assays”), approximately 10,000 cells of each strain were spotted side-by-side in the first position, and five-fold serial dilutions were spotted on adjacent positions, on plates with or without indicated stress conditions. Plates were incubated at 30°C and images were recorded daily for up to three days.

### Preparation of yeast lysate and Immunoprecipitation (IP)

Yeast cultures (30-40 mL) were grown in appropriate liquid medium to an OD_595nm_ of 0.5-0.7. Cultures were then treated with stress conditions, if appropriate, and harvested by centrifugation, followed by a wash with IP buffer (50 mM Tris-HCl, pH 8, 150 mM NaCl, 2.5 mg/mL *N*-ethylmaleimide, 0.1% Nonidet P-40 (NP40), 1X yeast protease inhibitor cocktail (BioShop), and 1 mM phenylmethylsulfonyl fluoride (PMSF)). Cells were then broken up with glass beads in IP buffer containing 0.1 mM dithiothreitol for 30 min at 4°C with a 5 min ice incubation halfway through. The lysed materials were isolated from the beads and separated from cell debris by centrifugation three times at 14,000 *g* for 5 min. The soluble samples were either analyzed by immunoblot or used for IP. If proceeding with immunoblot experiments, the samples were diluted with an equal volume of 2X SDS-PAGE sample buffer (4% sodium dodecyl sulfate, SDS; 20% glycerol; Bromophenol Blue; 10% 2-Mercaptoethanol; and 140 mM Tris-HCl pH 8) and boiled for 4 min prior to analysis by the indicated immunoblots. For IP experiments, 50 μL yeast lysate was retained as input sample, and the remainder was incubated for 2 h with anti-HA agarose beads at 4°C. IPs were then washed three times with ice-cold IP buffer containing 0.1% NP40, and twice with IP buffer alone. Beads were then resuspended with 2X SDS-PAGE sample buffer and boiled for 4 min prior to analysis by the indicated immunoblots. Quantification of Cst6 sumoylation levels were obtained using ImageJ program. First, on the SUMO immunoblot, intensity of signals below the non-specific band were quantified, then these were normalized to the intensity of corresponding HA IP signals. Finally the normalized values plotted relative to the control (first) sample. Statistically significant differences, calculated using Student’s *t*-test (*P* < 0.05), are denoted with an asterisk on graphs.

### Denatured immunoprecipitation (IP)

Denatured IP samples were prepared using a method previously described by Sri Theivakadadcham *et al.,* (2019). Briefly, 50 mL of yeast cultures were grown in rich medium, harvested by centrifugation, and washed with 20% Trichloroacetic acid (TCA). Lysed, precipitated and washed materials were resuspended with modified SDS buffer (60 mM Tris pH6.7, 5% 2-mercaptoethanol, 1% SDS, and few drops of bromophenol blue) prior to boiling for 5 min. These samples were then centrifuged to remove insoluble material. The supernatant was separated into a new tube and an aliquot (40 μL) of this supernatant was retained as input sample and the remainder was diluted with denaturing IP buffer (50 mM Tris pH 7.4, 150 mM NaCl, and 0.5% NP40) containing 0.5 mg/mL of bovine serum albumin. Diluted samples were then incubated overnight with anti-HA agarose beads at 4°C. The next day, IP samples were washed with ice-cold denaturing IP buffer. Beads were then resuspended with 2X SDS-PAGE sample buffer and boiled for 3 min prior to analysis by the indicated immunoblots.

### Chromatin immunoprecipitation (ChIP)

ChIP samples were prepared using a method previously described by Sri Theivakadadcham *et al.,* (2019). 50 mL yeast cultures were grown in rich medium and ethanol stress was induced by adding absolute ethanol to a final concentration of 5% for the indicated time points. Cells were then cross-linked with 1.1% of formaldehyde for 20 min, followed by 5 min of quenching with 282 mM of glycine and occasional mixing. Cells were harvested by centrifugation at 3000 *g* for 5 min and washed twice with ice-cold TBS (20 mM Tris-HCl, pH 7.5 and 150 mM NaCl). Washed samples were resuspended in cold ChIP buffer (50 mM HEPES-KOH, pH 7.5, 150 mM NaCl, 1 mM EDTA, 1% Triton X-100, and 0.1% sodium deoxycholate and 0.1% SDS) containing 1 mM PMSF and 1X yeast protease inhibitor cocktail. Cells were disrupted and lysed with glass beads using a mini bead beater. The homogenized sample was isolated from glass beads and sonicated to obtain an average DNA fragment size of ~500 bp in length. Samples were then centrifuged for 5 min at 14, 000 *g*, and the supernatant was transferred to a new tube. Then, NaCl was added to the supernatant to a final concentration of 225 mM prior to overnight incubation with 15 μL of magnetic Protein G beads (Dynabeads, Thermo Fisher Scientific) pre-bound with 2 μg rabbit anti-HA (Novus). On the next day, IPs were washed with four different buffers for 4 min each in the following order: (1) ChIP buffer with 275 mM NaCl; (2) ChIP buffer with 500 mM NaCl; (3) 10 mM Tris-HCl, pH 8, 0.25 M LiCl, 1 mM EDTA, 0.5% NP-40 and 0.5% sodium deoxycholate; (4) Tris-EDTA buffer (10 mM Tris-HCl, pH 8 and 1 mM EDTA).Washed beads were then incubated in ChIP elution buffer (50 mM Tris-HCl, pH 7.5, 10 mM EDTA and 1% SDS) for 20 min at 65°C. HA bound materials were isolated on a magnet stand and were then treated first with RNase for 30 min at 37°C and then again with proteinase K for 1 hour at 42°C. To reverse crosslinking, samples were incubated overnight at 65°C. Next day, DNA was purified and recovered using the GeneJet Gel Extraction Kit (Thermo Fisher). Purified DNA was used for quantitative PCR using primers listed in Table S2. Results from qPCR were normalized to an internal control gene, an untranscribed region of Chromosome V. Error bars represent the standard deviation of three replicates and the asterisks indicate a significant difference with *P*-values less than 0.05 obtained from a Student’s t-test.

### Isolation of RNA and reverse transcription (RT)

Yeast cultures (10 mL) were grown in rich medium and ethanol stress was induced by adding absolute ethanol to a final concentration of 5% for the indicated time points. RNA was isolated as previously reported (Amberg et al. 2006). Briefly, samples were harvested by centrifugation at 3000 *g* for 3 min and washed twice with chilled AE buffer (50 mM sodium acetate, pH 5.2 & 10 mM EDT A, pH 8.0) made with DEPC-treated RNase-free water. The pellet was resuspended with ice cold AE buffer and mixed with SDS, and phenol, pH 4.5. Samples were lysed using a freeze/thaw/vortex cycle twice. First the samples were chilled in a dry ice/ethanol bath slurry mixture (freezing step) for 5 min and then transferred to a 65°C water bath (thawing step) for 5 min, followed by vortex for 30 s. Samples were kept in the dry ice/ethanol bath for a final 5 min before centrifuging at 14, 000 *g* for 7 min at room temperature. RNA was precipitated using the aqueous layer. The isolated RNA was subjected to DNAse treatment followed by cDNA synthesis using the iScript reverse transcriptase (Bio-Rad). Quantitative PCR was performed using primers listed in Table S2. mRNA levels were normalized to an internal loading control, 25S rRNA. Error bar represents the standard deviation of three replicates and the asterisks indicate a significant difference with *P*-values less than 0.05 obtained from a Student’s t-test.

## ACKNOWLEDGEMENTS

This work was supported by a grant from the Canadian Institutes of Health Research (CIHR; grant number MOP-142282) to E.R.

## SUPPLEMENTARY FIGURE LEGENDS

**Figure S1: (A) Presence of HA tag does not affect growth in rich media containing glucose as the carbon source**. Cell growth of Cst6-6HA and parent strains was assessed on glucose-containing, rich medium plates under non-stress and indicated ethanol stress conditions by spot assay. Plates were incubated two days at 30°C. **(B) Presence of HA tag does not affect growth in rich media containing ethanol as the carbon source.** Growth fitness was assessed by spot assay on media containing either 1% or 3% ethanol as the sole carbon source, in the absence or presence of oxidative stress (1 mM H2O2). Plates were incubated for two days at 30°C. **(C) Cst6 sumoylation is detected in samples prepared with NEM and with intact lysine residues 139, 461 and 547.** Cst6-6HA WT and Cst6-6HA MT were IPed using HA-conjugated beads from cell lysates prepared with or without NEM and analyzed by HA and SUMO immunoblots. The experiment was also performed in the presence and absence of 5% ethanol stress, where indicated. Red asterisk (*) indicates the SUMO modified form of Cst6 on the HA blot.

**Figure S2: Global levels of sumoylation are influenced by carbon source and ethanol stress. (A)** Yeast cells show significantly higher levels of sumoylation when grown in media containing glucose compared to ethanol. Cells were grown in SC or rich media containing either 2% glucose (Glu) or 1% ethanol (EtOH) as a carbon source, then treated with or without a stressor (100 mM H2O2 for ethanol-containing medium, or 5% ethanol for glucose-containing medium) for 20 min. Lysates were prepared and analyzed by SUMO and GAPDH immunoblots. **(B)** Global sumoylation during ethanol stress is dose dependent. Lysates from cells treated with the indicated concentrations of ethanol for 20 min were analyzed by SUMO and GAPDH immunoblots. **(C)** Quantification of sumoylation levels for *B.* SUMO signals were normalized to corresponding GAPDH signals, and the results were plotted relative to the first sample. Error bars represent standard deviation of three replicates.

