## Supplementary figures and images for "Ethanol stress stimulates sumoylation of transcription factor Cst6 which restricts expression of its target genes"

### Figure S1

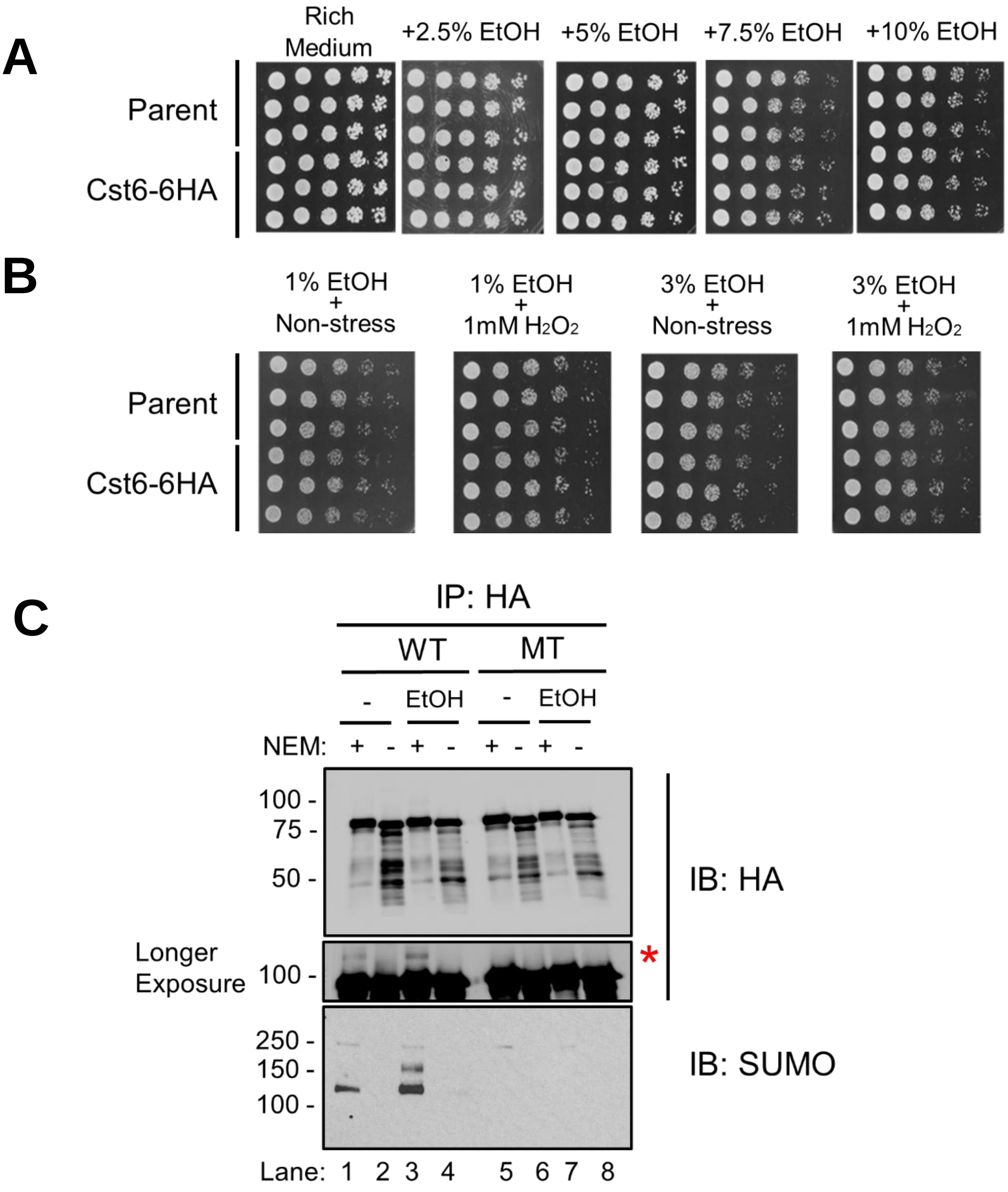

### Figure S2

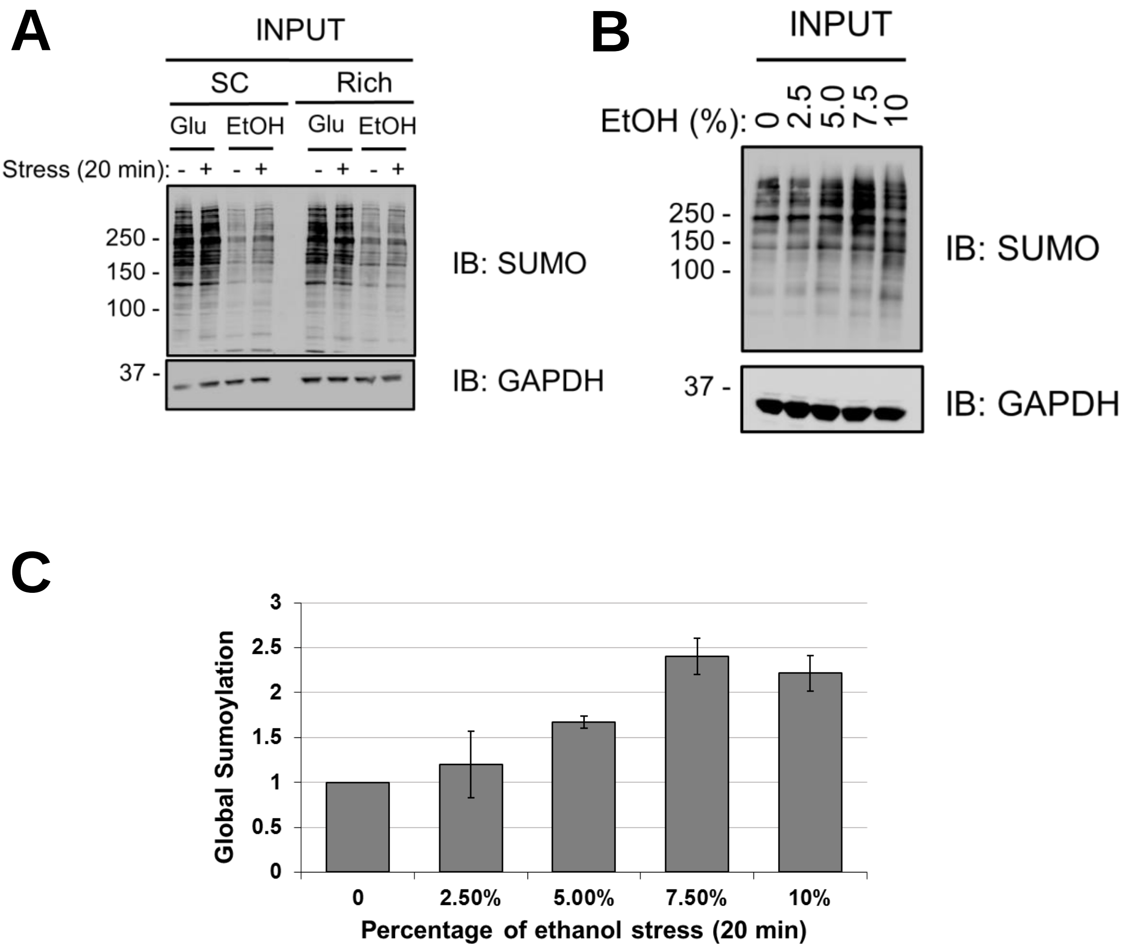
