## Supplementary material for "Ethanol stress stimulates sumoylation of transcription factor Cst6 which restricts expression of its target genes": Table S1

**Supplementary Table S1. Yeast strains used in this study**

| Strain | Parental | Genotype |
|---|---|---|
| ***Background Strain*** | | |
| W303a | | *MAT a ura3-52 trp1Δ2 leu2-3_112 his3-11 ade2-1 can1-100* |
| ***Derived Strains*** | | |
| YVS003C | W303a | *SKO1-6HA::kl TRP1* |
| YVS004A1 | W303a | *CST6-6HA::kl TRP1* |
| YVS064G | W303a | *cst6-K139R-6HA:: kl TRP1* |
| YVS060C | W303a | *cst6-K461R-6HA:: kl TRP1* |
| YVS061D | W303a | *cst6-K547R-6HA:: kl TRP1* |
| YVS063C | W303a | *cst6-K461,547R-6HA:: kl TRP1* |
| YVS065C | W303a | *cst6-K139,461,547R-6HA:: kl TRP1* |
| YVS062A | W303a | *cst6Δ::kanMX* |
