## Supplementary material for "Ethanol stress stimulates sumoylation of transcription factor Cst6 which restricts expression of its target genes": Table S2

**Supplementary Table S2. Oligonucleotide primers used in this study.**

| **Gene** | **Oligonucleotide sequence(s)** |
|---|---|
| ***Primers for quantitative PCR analysis of ChIP samples*** | |
| *NCE103* promoter | Forward: 5′-CGTTCGTTCCCGTTTCGATG-3′ |
| | Reverse: 5′-CGGCTTGATAAGCGTCATGG-3′ |
| *PYC1* promoter | Forward: 5′-CTTAGACAAGCGCCAGTTGC-3′ |
| | Reverse: 5′-CACTGCGGAAAAGGCCAAAA-3′ |
| *YAP6* promoter | Forward: 5′-CTCCCACAGTCACGATGAGC-3′ |
| | Reverse: 5′-TAGTCTCAACGGCTGTGCAT-3′ |
| *RPS3* promoter | Forward: 5′-GCAGGCATAAGATGCATAGCG-3′ |
| | Reverse: 5′-TCAAAAGCATCATGCGGCG-3′ |
| *ACC1* promoter | Forward: 5′-GCATGCATCGACGTCACAGT-3′ |
| | Reverse: 5′-CTAGTCGCAGTCGTGGTGAC-3′ |
| *PMA1* promoter | Forward: 5′-CAATTATGACCGGTGACGAAAC-3′ |
| | Reverse: 5′-AATCGAAACTAATGGAGGGGAG-3′ |
| *CHRV* untranscribed region | Forward: 5′-CATTATCCGTAACGCCACTTT-3′ |
| | Reverse: 5′-CGATCTTAGTTCCAATGGTGAAA-3′ |
| ***Primers for quantitative RT-PCR analysis*** | |
| *NCE103* | Forward: 5′-ACCCTACTGTGCAAACTGCT-3′ |
| | Reverse: 5′-GCAGTAGACCGTCCTCTACG-3′ |
| *ROX1* | Forward: 5′-GTCCACAACTACCCCTACGC-3′ |
| | Reverse: 5′-TAGCGGTGACCTCAGTGTTG-3′ |
| *ACC1* | Forward: 5′-TCCAGAAGATGTCGAAGCCG-3′ |
| | Reverse: 5′-TCAAGGCACGCAATGGTACT-3′ |
| *RPS3* | Forward: 5′-TCTGGTCAACCAGTCAACGAC-3′ |
| | Reverse: 5′-AGCCTTTGGACCAGTTCTGC-3′ |
| *YAP6* | Forward: 5′-GGACTTCCGAACCAGAGCAT-3′ |
| | Reverse: 5′-GGTATTGCGAGATGGGAGGG-3′ |
| *HAP4* | Forward: 5′-GTATTGACGGTAGTGCCGGT-3′ |
| | Reverse: 5′-TGGTGGCAGTTGCATCATTG-3′ |
| *25S* | Forward: 5′- TCTAGCATTCAAGGTCCCATTC-3′ |
| | Reverse: 5′- CCCTTAGGACATCTGCGTTATC-3′ |
